# Gypenosides modulate NCX calcium flux, insulin secretion and cytoprotection in BRIN-BD11 pancreatic β-cells

**DOI:** 10.1101/2021.01.29.428823

**Authors:** Chinmai Patibandla, Xinhua Shu, Angus M Shaw, Sharron Dolan, Steven Patterson

**Author notes:** **Corresponding author**: Steven Patterson PhD, Department of Biological and Biomedical Sciences, School of Health and Life Sciences, Glasgow Caledonian University, Cowcaddens road, Glasgow, G4 0BA.

## Abstract

Gypenosides are saponins extracted from the plant *Gynostemma pentaphyllum*, suggested to have antidiabetic and anti-obesity potential. However, its mechanism of action is not fully understood. The present study aimed to investigate the cytoprotective and insulin stimulatory effects of gypenosides using the rat BRIN-BD11 β-cell line. Gypenosides provided a significant cytoprotective effect against palmitate-, peroxide- and cytokine-induced cytotoxicity, with upregulation of antioxidant genes *Nrf2*, *Cat*, *Sod1,* and *Gpx1*. Acutely, gypenosides enhanced intracellular calcium ([Ca^2+^]_i_) and insulin secretion in a dose-dependent manner. The presence of the sodium/calcium exchanger (NCX) reverse mode inhibitor SN-6 blocked the gypenosides mediated increase in [Ca^2+^]_I_ but not the insulin secretion. These findings indicate that gypenosides may enhance [Ca^2+^]_i_ by activating the reverse mode of NCX channels and a possible calcium-independent mechanism involved in their insulin secretion. Gypenosides also upregulate the antioxidant gene expression and protect against oxidative stress and lipotoxicity, providing the rationale for their observed antidiabetic actions.

## Introduction

Type 2 diabetes mellitus (T2DM) is a metabolic disorder characterised by reduced peripheral insulin sensitivity, β-cell stress, and dysfunction. This ultimately leads to reduced functional β-cell mass, increasing glucose intolerance, and resultant hyperglycemia. Glucose-stimulated insulin secretion (GSIS) requires glucose metabolism and mitochondrial ATP production. The increase in the ATP/ADP ratio causes closure of ATP-sensitive K^+^ channels, membrane depolarisation, the opening of voltage-gated Ca^2+^ channels, and increasing intracellular Ca^2+^ levels, which drives insulin exocytosis (Fu et al., 2013). GSIS also results in the production of reactive oxygen species (ROS) as a by-product during mitochondrial metabolism of glucose in the β-cell. However, these ROS are rapidly converted to less toxic molecules by antioxidant enzymes (Brookes et al., 2004). Prolonged insulin hypersecretion by β-cells to compensate for peripheral insulin resistance causes intracellular ROS accumulation in β-cells. Due to the very low expression of antioxidant enzymes in the β-cells, combined with accumulating ROS, β-cells are susceptible to endoplasmic reticulum stress, initiating proinflammatory responses and eventually causing β-cell apoptosis (Donath et al., 2009). In *in vivo* and *in vitro* studies, activation of *Nrf2* (a master regulator of antioxidant pathways) triggered β-cell self-repair and protected against oxidative stress (Abebe et al., 2017; Bhakkiyalakshmi et al., 2014). Thus, any insulin secretagogue with β-cell protective effects against ROS may have therapeutic potential in T2DM.

GYP are dammarane type triterpene glycosides extracted from *Gynostemma pentaphyllum* (GP), which structurally resembles ginsenosides of *Panax ginseng* (Bai et al., 2010). GYP has previously shown to have anti-hyperglycaemic and hypo-lipidaemic properties in Zucker fatty rats (Megalli et al., 2006). The herbal extract of GP also reduced hepatic glucose output and enhanced insulin secretion in diabetic Goto-Kakizaki rats (Lokman et al., 2015; Yassin et al., 2011). In T2DM patients, GP tea consumption enhanced insulin sensitivity when taken on its own or in combination with sulfonylureas (Huyen et al., 2012, 2013). Furthermore, in isolated pancreatic islets from healthy Wistar rats and spontaneously diabetic Goto-Kakizaki rats, Phanoside (a GYP extracted from GP) enhanced insulin secretion at both low (3.3mM) and high (16.7mM) glucose concentrations (Hoa et al., 2007).

In β-cells, intracellular calcium plays a significant role as a secondary messenger promoting insulin granule docking and fusion with the plasma membrane and release of insulin by exocytosis (Newsholme et al., 2012). There are many calcium channels expressed in β-cells, including L-type, T-type, P/Q type, store-operated calcium channels (SOCC), and sodium-calcium exchanger (NCX). Previous attempts to find the calcium channels involved in GYP-induced insulin secretion were inconclusive (Lokman et al., 2015). In streptozotocin-induced diabetic rats, GYP showed antidiabetic effect by stimulating insulin secretion, reducing glucose and lipid levels, enhancing *Nrf2* and its associated antioxidant gene expression (Gao et al., 2016). A similar *Nrf2* mediated protective effect of GYP was also reported in PC12 cells (neural differentiation cell model) and ARPE19 (retinal pigmental epithelial cells) cells (Alhasani et al., 2018; Meng et al., 2014).

The current study focused on elucidating the effects of GYP on insulin secretion and β-cell function, Ca^2+^ signaling, and cytoprotection in the rat clonal BRIN-BD11 β-cell model.

## Methods

### Gypenoside extract preparation

Gypenosides were purchased from Xi’an Jiatian Biotech Co. Ltd, China (purity 98%). Gypenosides were dissolved in absolute ethanol (25mg/ml) by continuous shaking at room temperature overnight. The extract was sterile filtered through a 0.2μm filter and stored at −20°C until use.

### Solutions and Chemicals

Krebs Ringer bicarbonate buffer (KRBB) was composed of (mmol/L): 115 NaCl, 4.7 KCl, 1.2 KH_2_PO_4_, 1.2 MgSO_4_, 1.28 CaCl_2_, 20 HEPES, 24NaHCO_3_ and 0.1% (w/v) bovine serum albumin (pH 7.4). Thapsigargin (1138), SKF96365 (1147), and SN-6 (2184) were purchased from Tocris, UK. All other chemicals were from Sigma Aldrich (Poole, UK) unless indicated. For studies with palmitic acid, the palmitic acid was dissolved in 50% ethanol at 70°C and conjugated to 10% (w/v) fatty acid-free BSA in RPMI for 1h at 37°C with constant stirring. The respective palmitic acid stock was diluted 1:10 to give the final concentrations (125μM or 250μM) of palmitate and 1% (w/v) BSA. Final concentrations of ethanol in cell culture medium never exceeded 0.01% (v/v).

### BRIN-BD11 cell culture and viability testing

BRIN-BD11 cells (a kind gift from Prof. Peter Flatt, Ulster University, UK) were cultured in RPMI 1640 medium with 2mM L-Glutamine (Lonza, Belgium), supplemented with 10% (v/v) fetal bovine serum (FBS), 50U/ml penicillin/streptomycin and maintained at 37°C with 5% CO_2_ and 95% air. Cells were trypsinised and sub-cultured at 1:5 dilutions when 80-90% confluence was reached. Passages between 25-40 were used for the experiments. To determine the effects of GYP on cell viability, BRIN-BD11 cells were seeded on 96 well plates at 1×10^4^ cells/ well. After overnight culture, cells were incubated with test reagents as indicated in the figures for 6 – 72h and cell viability assessed using MTT assay as described previously (Vasu et al., 2014).

### Insulin Secretion studies

Cells were seeded onto a 12 well tissue culture plate (1.5 x 10^5^ cells/ well) and was incubated overnight in RPMI complete growth medium. For insulin release study, media was removed and cells washed with phosphate-buffered saline (PBS). Cells were preincubated for 40 min in KRBB containing low glucose (1.1mM), prior to replacement with low (1.1 mM) or high glucose (16.7mM) KRBB with or without GYP (1ml/well) and incubated for 1h at 37°C. After incubation, test buffer from each well was collected, and insulin levels were estimated using a total rat insulin ELISA kit (Merck Millipore, UK) according to the manufacturer’s instructions. An Epoch (BioTek, UK) microplate spectrophotometer was used to read the absorbance at 450nm and 590nm.

### Calcium Imaging

BRIN-BD11 cells were seeded onto glass coverslips (1.0 x 10^5^ cells/ coverslip) and allowed to attach overnight in RPMI 1640 complete media. Before use, cells were incubated for 45 min with 2μM FURA-2AM (Tocris, UK) in KRBB containing 1.1 mM glucose at 37°C. Coverslips were rinsed with PBS and mounted onto an RC-21BRW closed bath imaging chamber (Warner Instruments) with a P-2 platform. Cells perfused at 1ml/min in low glucose (1.1mM) KRBB for 30 min before assessing intracellular calcium. A Nikon Eclipse TE2000-U microscope fitted with Photometrics Cool SNAP^™^ HQ Camera was used to acquire images. The Fura-2 340/380 ratio was calculated, and graphs plotted using MetaFluor fluorescence ratio imaging software.

### Quantitative real-time PCR

Total RNA was extracted from GYP pre-treated BRIN-BD11 cells using NucleoSpin® RNA kit (Macherey-Nagel, UK) according to the manufacturer’s protocol. From total RNA, cDNA was synthesised using a High-Capacity cDNA reverse transcription kit (Applied Biosystems, UK). Changes in expression of mRNA levels for genes of interest and reference genes were measured by quantitative real-time PCR using 5X HOT FIREPol® EvaGreen® qPCR Mix Plus (no ROX) (Solis BioDyne, Estonia). Bio-rad CFX96™ Real-Time PCR detection was used for amplification and detection. Cycling conditions used were, 95°C for 12min followed by 40 cycles of denaturation (95°C for 15s), annealing (60°C for 20s) and extension (72°C for 20s). At the end of each experiment, melting curve analysis was done to analyse primer specificity.

### Western blot

Cells were lysed with RIPA buffer (150mM NaCl, 0.1% Triton X-100, 0.5% Sodium deoxycholate, 0.5% SDS and 50mM Tris-HCl, pH8.0), sonicated and centrifuged to separate any cell debris. Protein concentration in the supernatant was measured using a DC™ Protein assay kit (BIO-RAD, UK) according to the manufacturer’s specifications. Proteins were separated by SDS-PAGE and transferred to nitrocellulose membrane using iBlot® blotting system (Thermo Scientific, UK). Membranes were blocked with 5% (w/v) BSA in Tris-buffered saline containing Tween20 for 1h at room temperature and incubated with primary antibodies overnight at 4°C. Primary antibodies and dilutions used were: Anti-Pcsk1 (1:1000) (GTX113797S, Genetex), Anti-Pcsk2 (2μg/ml) (MAB6018-SP, Novus Biologicals), Anti-Pdx1 (1μg/ml) (AF2517, R&D Systems) and Anti-Actin-beta (1:1000) (ab-1801, Abcam UK). Blots were incubated with IRDye® conjugated specific secondary antibodies (1:5000) for 1h at room temperature. Signal was detected using Odyssey® Fc imaging system (LI-COR, UK) and analysed using Image Studio™ software.

### Data Analysis

Results were presented as mean ± S.E.M. Data was analysed using Graphpad PRISM® software (ver 6.01) and unpaired t-test (parametric) for comparing two groups or one-way ANOVA for comparing more than 2 groups with a significance threshold of p<0.05.

## Results

### GYP treatment dose-dependently alters BRIN-BD11 cell viability

Concentration-dependent long term (24-72h) effects of GYP on the viability of BRIN-BD11 cells are shown in figure 1. A higher concentration of GYP (100μg/ml) reduced cell viability over 24-72h treatment (P<0.0001). GYP at a 50μg/ml concentration over 24h treatment showed a slight but significant increase in viability (P<0.05). Although at 48h, 50μg/ml showed no significant effect, longer-term treatment (72h) significantly reduced the viability of the cells (P<0.0001). Low concentrations of GYP (25-6.25μg/ml) had no significant effect until 48h treatment, where 25 and 12.5μg/ml GYP treatment significantly enhanced cell viability by 1.17- and 1.33-fold (P<0.05 & P<0.01, respectively). At 72h treatment, 1.9, 1.7- and 1.6-fold increases in viability were observed when treated with 25, 12.5, and 6.25 μg/ml GYP, respectively (P<0.0001). As a low concentration of 12.5μg/ml GYP enhanced cell viability significantly at 48 and 72h (1.33 and 1.6-fold respectively), this concentration was used for further testing of GYP’s potential protective effects against cytotoxic concentrations of palmitate, peroxide, and a cytokine cocktail. Acute (1-6 h) exposure of cells to all GYP concentrations tested did not reduce cell viability (data not shown).

**Figure 1:**
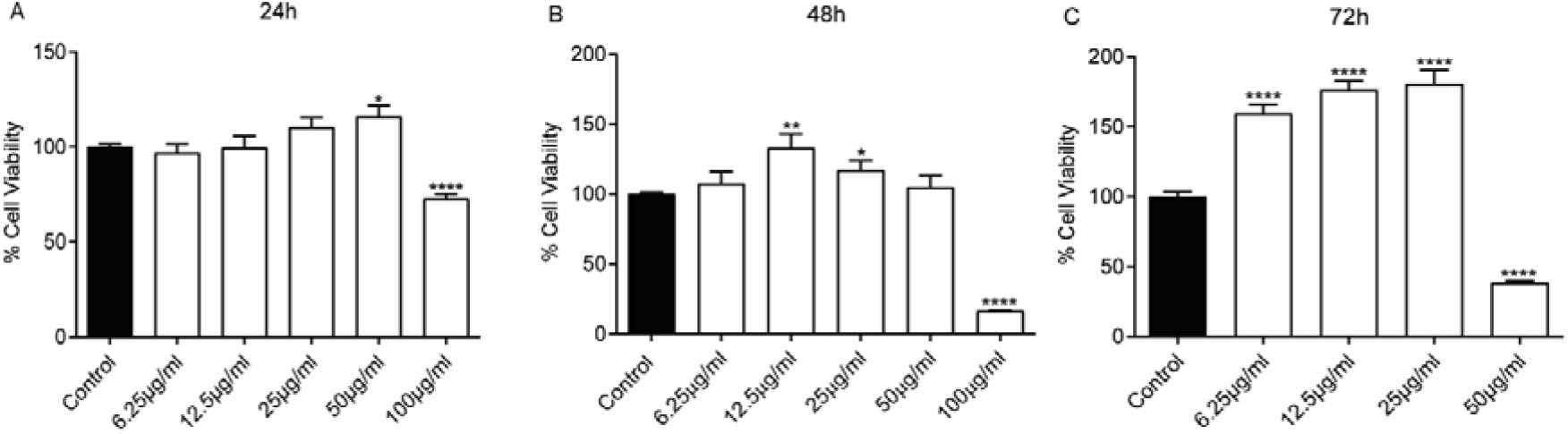
Effects of GYP addition to BRIN-BD11 cell culture medium at concentrations between 6.25–100 μg/ml on cell viability following 24h (A), 48 h (B), and 72h (C) treatment. Plotted as % change in cell viability compared to control. Values represent mean ±S.E.M. from three different experiments (n=4). The student’s t-test was used for statistical analysis. *P<0.05; **P<0.01; ****P<0.0001, compared to control (no GYP addition).

### GYP has cytoprotective effects against palmitate-, peroxide- and cytokine-induced toxicity

Effects of low concentrations of GYP (10, 12.5, and 15μg/ml) against 125 and 250μM palmitate-induced toxicity are shown in figure 2A. Palmitate treatment for 24h at a concentration of 125 and 250 μM significantly reduced cell viability (P<0.0001) by 50% and 70%, respectively. Low concentrations of GYP (10 & 12.5μg/ml) in the presence of 125 μM palmitate significantly protected cells from the detrimental actions of palmitate (P<0.001 and P<0.0001, respectively). Treatment of BRIN-BD11 cells with GYP (10, 12.5, and 15 μg/ml) along with higher concentrations (250 μM) of palmitate also significantly reduced the decline in cell viability (P<0.001-0.0001).

**Figure 2:**
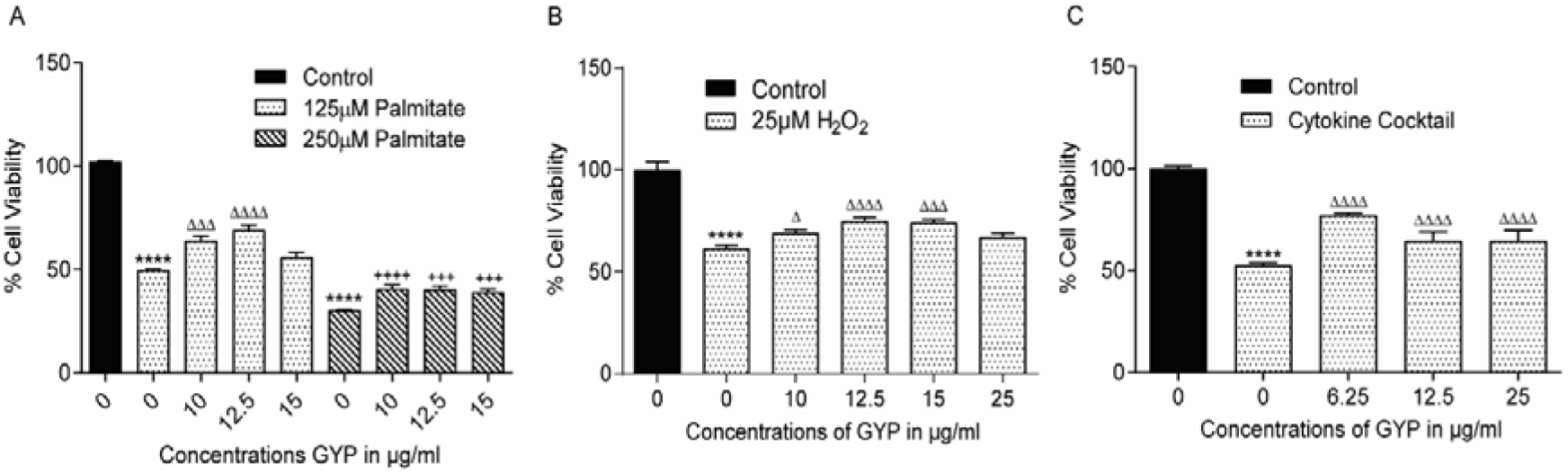
Protective effects of GYP against detrimental effects of palmitate (A), peroxide (B), and cytokine cocktail (C) on BRIN-BD11 cell viability. Cells were cultured with palmitate and GYP for 24 h prior to cell viability measurement by MTT assay. For H_2_O_2,_ cells were exposed to treatment for 6h alone or in combination with GYP. For cytokine treatment, cells were exposed to a cytokine cocktail containing IL-1β (50U), TNF-α (1000U), and IFN-γ (1000U) for 24 h either alone or in combination with GYP. Values represent mean ±S.E.M. from three different experiments (n=4). The student’s t-test was used for statistical analysis. ****P<0.0001 compared to control (black bar); ^Δ^P<0.05, ^ΔΔΔ^P<0.001, ^ΔΔΔΔ^P<0.0001 compared to 125 □M palmitate, H_2_O_2_ or cytokine cocktail treatment alone.; ^+++^P<0.001; ^++++^P<0.0001 compared to 250 □M palmitate alone.

Peroxide treatment for 6h reduced (P<0.0001) cell viability compared to untreated cells (Figure 2B). The addition of GYP (10, 12.5, and 15μg/ml) protected the cells (P<0.05-P<0.0001) against the detrimental effects of peroxide.

Protective effects of GYP over 24h against inflammatory cytokine cocktail-induced toxicity is shown in figure 2C. The cytokine cocktail containing IL-1β (50U), TNF-α (1000U), and IFN-γ (1000U) reduced cell viability by 61% (P<0.0001) over 24h. GYP co-treatment (6.25, 12.5, and 25μg/ml) protected the cells against cytokine-induced toxicity (P<0.0001), with increased viability of 1.1, 1.2, and 1.2-fold, respectively when compared to cytokine cocktail treatment alone.

### GYP promote insulin secretion from BRIN-BD11 cells irrespective of glucose concentration

BRIN-BD11 cells were incubated in KRBB containing either low (1.1mM) glucose or high (16.7mM) glucose with different concentrations of GYP (6.25, 12.5, 25, 50, and 100μg/ml) for 1h, and insulin was measured by ELISA (figure 3). High glucose (16.7mM) significantly increased insulin secretion (P<0.05) compared to low glucose. Low concentrations of GYP (6.25, 12.5, and 25μg/ml) had no insulin release effects at both low and high glucose. However, GYP at 50μg/ml increased insulin release 1.9-fold compared to 1.1mM glucose alone (P<0.01) but had no effect at high glucose. GYP at 100μg/ml enhanced insulin secretion 4.4-fold (P<0.0001) at 1.1mM glucose and 3-fold (P<0.001) at 16.7mM glucose compared to respective controls. This was similar to secretion in the presence of 10mM L-alanine with increases in insulin secretion of 5-(P<0.0001) and 3-(P<0.0001) fold at basal and high glucose, respectively.

**Figure 3:**
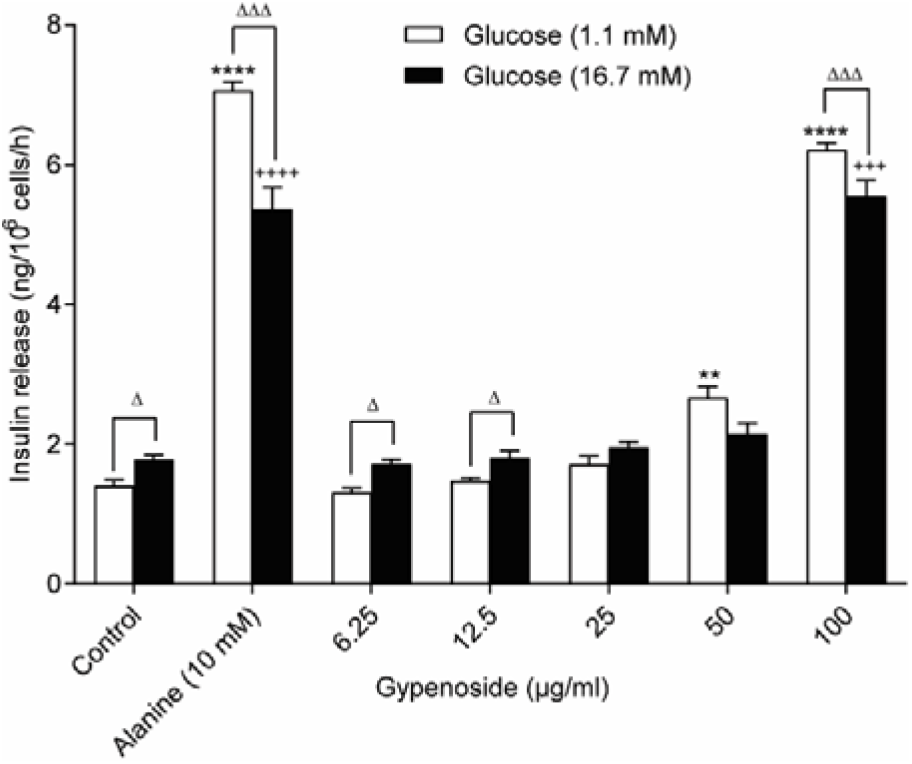
Concentration-dependent effects of gypenosides on insulin secretion from BRIN-BD11 cells. Insulin secretion measured in presence of GYP 6.25-100μg/ml at low (1.1mM) (white bars) or high (16.7mM) (black bars) glucose concentrations. Data plotted as concentration of insulin secreted in ng/10^6^ cells per h. Values plotted as mean ± S.E.M. from 4 independent experiments conducted in duplicate. **P<0.01, ****P<0.0001 compared to 1.1 mM glucose control; ^Δ^P<0.05, ^ΔΔΔ^P<0.001 compared to respective 1.1 mM glucose result; ^+++^P<0.001, ^++++^P<0.0001 compared to 16.7 mM glucose control.

### NCX channels are involved in GYP induced Ca2^+^ uptake but not in insulin secretion

Acutely, GYP (50 & 100μg/ml) enhances BRIN-BD11 cell [Ca^2+^]_i_ levels at both low (1.1mM) and high glucose (16.7mM) which is consistent with GYP ability to enhance insulin secretion (Figure 4A & 4B). To determine the effect of GYP on intracellular stores, ER stores were emptied by the addition of 300nM thapsigargin followed by 100μg/ml GYP treatment along with the thapsigargin. As expected, thapsigargin significantly increased [Ca^2+^]_C_ while the addition of GYP with thapsigargin further enhanced [Ca^2+^]_i_ levels, indicating GYP-induced extracellular calcium entry through plasma membrane-bound calcium channels (Figure 4C). Verapamil and mibefradil, L-type and T-type calcium channel blockers, respectively, were used to establish if GYP acted via these channels. GYP-induced Ca^2+^ uptake was unchanged in the presence of either 10μM verapamil or 10μM mibefradil (Figure 4D&4E). To determine whether store-operated calcium channels (SOCC) are involved in GYP action, BRIN-BD11 cells were incubated with 30μM SOCC blocker SKF96365 along with 100μg/ml GYP. The presence of SOCC blocker delayed the time to response for GYP but did not block the Ca^2+^ entry induced by GYP (Figure 4F). SN-6 is an NCX reverse mode inhibitor that specifically blocks calcium entry through NCX. Perfusion of BRIN-BD11 cells with GYP (100μg/ml) in the presence of 10μM SN-6 completely blocked Ca^2+^ entry (Figure 4G), indicating NCX reverse mode activity might be involved in GYP-induced calcium responses. To further investigate if NCX mediated Ca^2+^ entry is involved in Insulin secretory properties of GYP, Insulin secretion was measured with GYP treatment in the presence of SN-6. The presence of SN-6 did not alter the insulin secretion by GYP (Figure 4H) at low glucose concentration, indicating Ca^2+^ independent pathways might be involved in the GYP mechanism.

**Figure 4:**
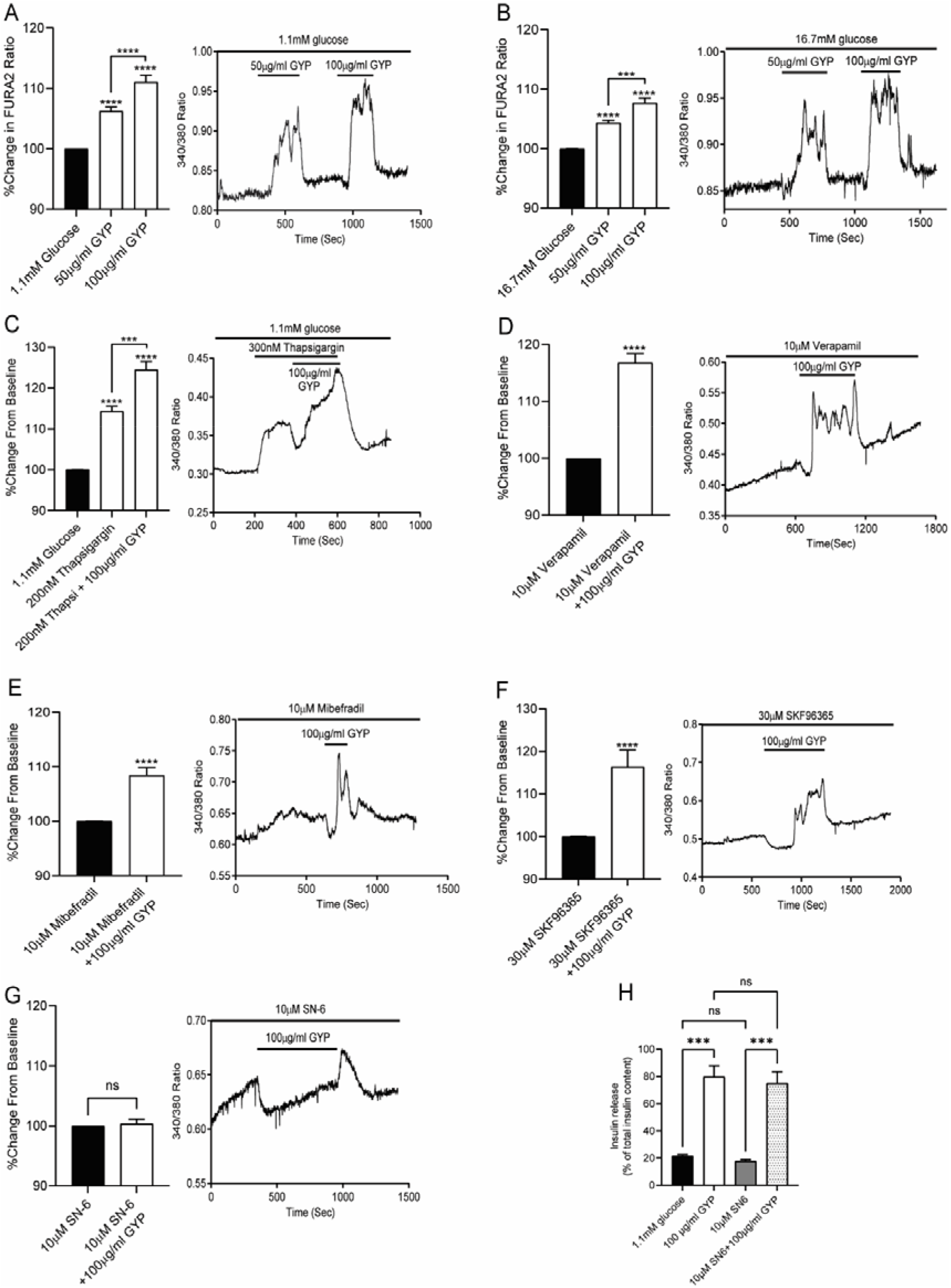
Effects of GYP on intracellular Ca^2+^ levels of BRIN-BD11 cells alone and in the presence of specific calcium channel modulators. Graphs showing the % change in FURA2 ratio (340/380) from baseline over time (left) and representative plot (right) when perfused with GYP in the presence of low (1.1 mM) (A) and high (16.7 mM) (B) glucose concentrations, Thapsigargin (C), L-type calcium channel blocker verapamil (D), T-type calcium channel blocker Mibefradil (E) SOCC blocker SKF96365 (F) and NCX channel reverse mode inhibitor SN-6 (G). Values plotted as mean responses of 3-15 responding cells from three independent experiments— Effects of SN-6 on GYP induced insulin secretion in 1.1mM glucose (H). Data plotted as % of insulin secreted from total content. Values plotted as mean ± S.E.M. from 3 independent experiments. ***P<0.001, ****P<0.0001.

### Treatment with GYP changes BRIN-BD11 gene expression

Following 24h treatment with GYP (12.5μg/ml), the expression of antioxidant genes *Sod1* (P<0.01) and *Cat* (P<0.05) were increased, while *Gpx1* expression was unchanged and *Ho1* expression was downregulated (P<0.0001) (Figure 5A). Interestingly, 72h treatment significantly increased expression of all antioxidant genes (*Sod1* (P<0.05), *Cat* (P<0.001), *Gpx1* (P<0.001), *Ho1* (P<0.001). However, no change in *Sod2* expression was observed at any time point.

**Figure 5:**
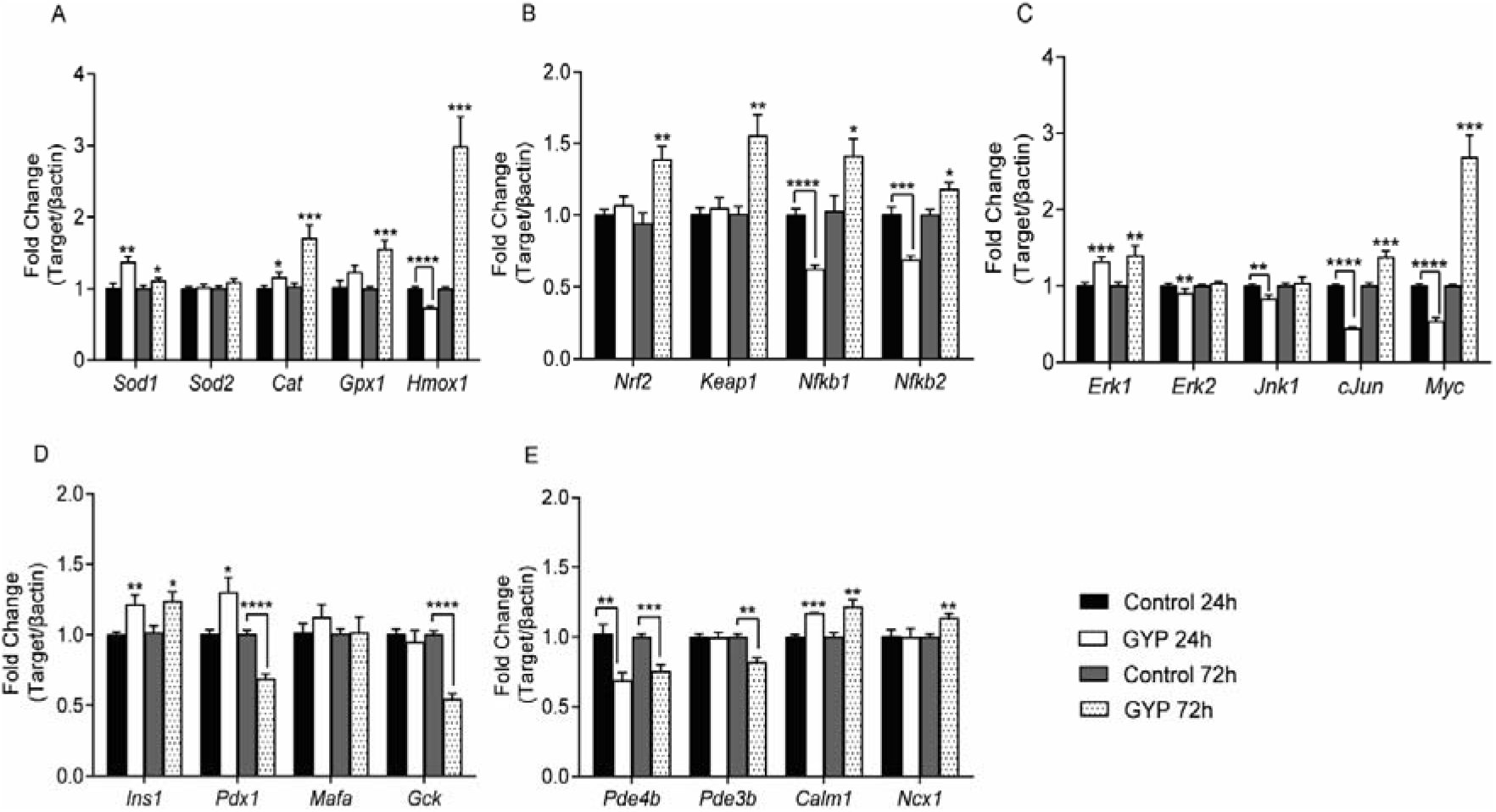
Effects of 24 and 72 h culture with GYP (12.5μg/ml) on the expression of antioxidant and βcell-specific genes in BRIN-BD11 cells. Data represent fold-change in mRNA levels compared to control/untreated BRIN-BD11 cell and normalised to β-actin expression. Values represent mean ±S.E.M. from three different experiments performed in duplicate. *P<0.05, **P<0.01, ***P<0.001, ****P<0.0001 compared to their respective 24h or 72h control.

As *Nrf2* regulates the expression of antioxidant genes, changes in expression of *Nrf2* along with its regulator, *Keap1*, were investigated following 24 & 72h GYP treatment. GYP treatment for 24h did not affect *Nrf2* and *Keap1* expression, while 72h treatment resulted in significant increases in both *Nrf2* and *Keap1* (P<0.01) (Figure 5B). *Nfkb* mediates cytokine-induced toxicity in β-cells, and as GYP treatment protected against cytokine-induced toxicity, changes in *Nfkb1* & *Nfkb2* were investigated. Both *Nfkb1* & *Nfkb2* expression was downregulated (P<0.0001) by GYP after 24h. However, unexpectedly a significant (P<0.05) increase in expression of both genes was observed following 72h GYP culture (Figure 5B).

Treatment with GYP for 24h enhanced *Erk1* expression (P<0.001) while *Erk2* (P<0.01), *Jnk1* (P<0.01), *cJun* (P<0.0001), and *cMyc* (P<0.0001) expression were downregulated (Figure 5C). At 72h, *Erk2* and *Jnk1* expression were unchanged, while *Erk1, cJun,* and *Myc* expression were upregulated (P<0.01, P<0.001 & P<0.001, respectively) (Figure 5C).

Expression of key β-cell genes, *Ins1* and *Pdx1,* increased significantly (P<0.01 and P<0.05, respectively) following 24h GYP treatment (Figure 5D), while glucokinase (*Gck*) and *MafA* expression were unchanged. Following 72h treatment of BRIN-BD11 with GYP *Ins1* expression was till raised (P<0.05), however, *Pdx1* and *Gck* expression were decreased (P<0.0001), and *MafA* expression remained unchanged (Figure 5D).

*Pde4b*, associated with cAMP degradation, was downregulated following 24h GYP treatment (P<0.01), although *Pde3b* expression was unchanged. Both *Pde4b* and *Pde3b* were significantly downregulated at 72h (P<0.001 and P<0.01, respectively) (Figure 5E). Calcium-associated calmodulin (*Calm1*) expression was upregulated (P<0.001) by 24h GYP treatment, while no change in NCX1 expression was observed. Both *Calm1* and NCX1 expression were upregulated following 72h GYP treatment (P<0.01) (Figure 5E).

### Long term treatment with GYP reduces the expression of *Pdx1* but not Prohormone convertases (*Pcsk1 & Pcsk2*)

Effects of 24h & 72h culture of BRIN-BD11 cells with GYP on specific changes in critical cellular proteins are shown in figures 6. As GYP treatment altered the expression of genes necessary for insulin production in the β-cells, protein levels of *Pdx1* (necessary for insulin gene transcription), *Pcsk1,* and *Pcsk2* (both necessary for post-translational modification of proinsulin to insulin) were determined by western blot analysis. At 24h, GYP (12.5μg/ml) had no significant effect on protein levels of *Pdx1*, while extended treatment for 72h significantly reduced *Pdx1* protein levels (Figure 6 A&B), consistent with changes seen at the mRNA levels shown in figure 5D. Expression of *Pcsk1* and *Pcsk2* were unchanged by GYP treatment.

**Figure 6:**
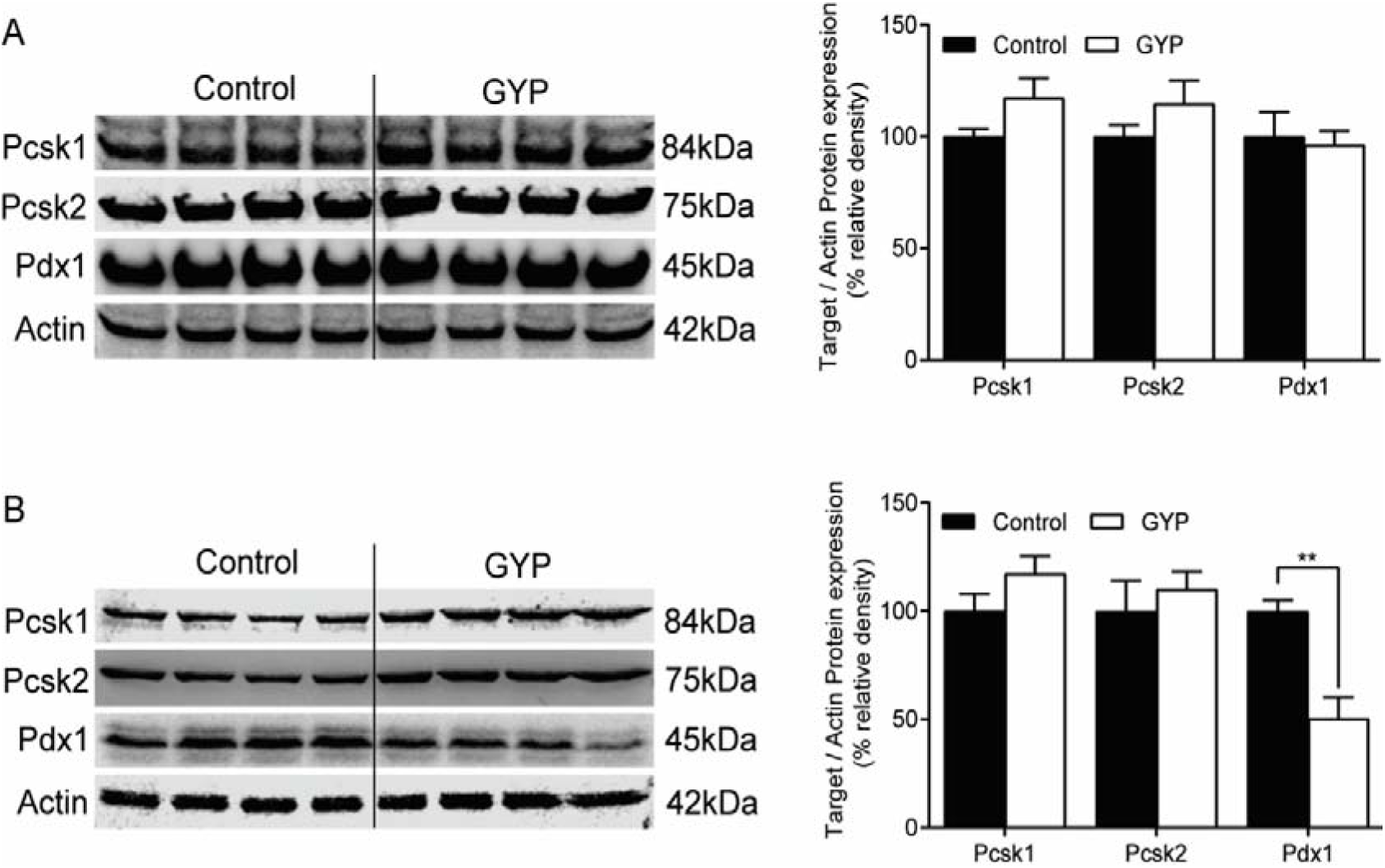
Effects of GYP (12.5μg/ml) treatment for 24 h (A, B) and 72 h (C, D) on the expression of Pdx1 and prohormone convertases 1 and 2 in BRIN-BD11 cells. Protein levels for each treatment were normalised to Actin expression, plotted as % change relative to control. Values represent mean ±S.E.M. from four different experiments. **P<0.01 compared to respective control.

## Discussion

Glucose-stimulated insulin secretion (GSIS) involves ATP-sensitive K^+^ channel closure, membrane depolarisation, and increased cytoplasmic calcium levels through the opening of voltage-dependent calcium channels (VDCC), including L-type, T-Type, and P/Q-type Ca^2+^ channels (Yang et al., 2006). The endoplasmic reticulum (ER) acts as an intracellular Ca^2+^ store, and its Ca^2+^ levels are maintained by Sarco-endoplasmic reticulum Ca^2+^-ATPase (SERCA) pumps. In GSIS, Ca^2+^ is released from ER stores into the cytoplasm along with extracellular Ca^2+^ influx. In the case of ER store Ca^2+^ depletion, store-operated calcium channels (SOCC) bound to the plasma membrane refill the ER and also enhance cytoplasmic Ca^2+^ (R. Wang et al., 2013). The rise in cytoplasmic Ca^2+^ ions, as a secondary messenger, is linked to many functions, including insulin exocytosis. In the current study, we report that GYP enhances [Ca^2+^]_i_ and insulin secretion in a concentration-dependent manner at low and high glucose concentrations. These results are consistent with previous results in isolated wild-type and diabetic Goto-kakizaki rat islets, where Phanoside (a Gypenoside) enhanced insulin secretion irrespective of glucose concentration (Hoa et al., 2007). The increase in [Ca^2+^]_i_ stimulated by GYP was unaffected by L-type calcium channel blocker verapamil, T-type calcium channel blocker mibefradil, and store-operated calcium channel blocker SKF96365. Thapsigargin is known to deplete the ER store calcium by blocking SERCA-induced calcium uptake into ER stores (Lytton et al., 1991). Thapsigargin induced ER depletion followed by GYP treatment enhanced [Ca^2+^]_i_ on top that caused by thapsigargin indicating extracellular calcium mobilisation mediated by plasma membrane-bound channels was key in GYP actions.

Plasma membrane Ca^2+^-ATPase (PMCA) and plasma membrane Na^+^/Ca^2+^ exchanger (NCX) extrude Ca^2+^ ions from the β-cell (Chen et al., 2003). NCX can work in a bidirectional manner; in forward mode, it expels Ca^2+^ and, in reverse mode, increases [Ca^2+^]_I_ (Philipson et al., 2002). It transports 3 Na^+^ ions for every Ca^2+^ ion. NCX exists in three isoforms NCX1, NCX2, and NCX3, encoded by genes *Slc8a1-3*. Rat pancreatic β-cells and β-cell models (RINm5F and BRIN-BD11) express two splice variants of NCX1 (NCX1.3 and NCX1.7) (F Van Eylen et al., 1997; Françoise Van Eylen et al., 2002). Benzyloxyphenyl derivatives like KB-R7943, SEA0400, and SN-6 are known for their NCX reverse mode inhibitory effects. Among the three, SN-6 is a potent and selective inhibitor of Ca^2+^ entry through NCX reverse mode (Iwamoto, 2004). We found that, in the presence of SN-6, GYP’s ability to stimulate and increase [Ca^2+^]_i_ was completely abolished, indicating GYP interacts with NCX reverse mode mediated Ca^2+^ entry. However, the Presence of SN-6 did not affect the GYP induced insulin secretion, indicating GYP might also promote calcium-independent insulin secretion pathways. It is previously reported that cAMP can potentiate insulin secretion independent of Ca^2+^ (Ämmälä et al., 1993; Kim et al., 2008), which could be the mechanism of GYP.

Reduced functional β-cell mass is characteristic of Type 1 diabetes and is also observed in Type 2 diabetes. Apoptosis is one of the significant causes for β-cell loss in T2DM and responsible for its progression (Butler et al., 2003). The contributing factors for β-cell apoptosis include inflammation involving an array of cytokines, oxidative stress caused by ROS/RNS, glucotoxicity due to prolonged hyperglycemia, and hyperlipidemia-induced lipotoxicity. Proinflammatory cytokines, specifically, interleukin 1-β (IL-1β), interferon γ (IFN-γ) and tumor necrosis factor-α (TNF-α), are linked to β-cell inflammation and apoptosis in Type1 diabetes mellitus (Donath et al., 2003). In T2DM, increased cytokine levels and immune cell infiltration have been observed in the islets, indicating an inflammatory response (Ehses et al., 2007). Although IL-1 β is produced in the β-cells in response to high glucose, the levels produced are not sufficient to cause apoptosis in purified rat, and human β-cells and combination with IFN-γ are necessary to promote apoptosis (Quan *et al.*, 2013). Exposure of rat and human β-cell models (BRIN-BD11 and 1.1B4) to cytokine cocktail of IL-1β, IFN-γ, and TNF-α, has previously been demonstrated to cause cellular apoptosis (Kiely et al., 2007; Vasu et al., 2014). Previous studies with GYP have shown anti-inflammatory properties in microglial cells (Cai *et al.*, 2016) and retinal pigment epithelial cells (Alhasani et al., 2018) by reducing cytokine levels. In chondrocytes, GYP was also able to protect against IL-1β mediated inflammation (Wan et al., 2017). Similar to these results, we have observed a protective effect of GYP in BRIN-BD11 cells against a decline in cell viability caused by a proinflammatory cytokine cocktail of IL-1β, IFN-γ, and TNF-α. Thus GYP may protect β-cells against inflammatory damage caused by metabolic disarray in obesity and diabetes.

Increased circulating free fatty acid (FFA) levels in obesity are one of the risk factors that are linked to the development and progression of type 2 diabetes. Chronic exposure to FFA due to substrate competition results in impaired glucose metabolism and favors FFA oxidation in the β-cell (Lupi et al., 2002). Acutely, FFA can stimulate insulin secretion through Gα_q/11_ coupled free fatty acid receptor 1 (*FFAR1/GPR40*) (Mancini et al., 2013). However, chronic exposure to elevated levels of FFA is linked to ER stress and β-cell apoptosis (Oh et al., 2018). Saturated FFA like palmitate is known to reduce GSIS and increase β-cell apoptosis in both human and rat models (Karaskov et al., 2006; Lupi et al., 2002). In primary hepatocytes, gypenoside treatment protected against palmitate-induced cell apoptosis (Müller et al., 2012). In our studies, a similar protective effect against palmitate in BRIN-BD11 cells was observed when GYP was added to the cells, further supporting the idea that GYP has broad-ranging beneficial cytoprotective effects in β-cells.

ROS (Superoxide O_2_^−^ and hydroxyl radical [HO^.^]) are by-products of the mitochondrial respiratory chain, and their production is linked to mitochondrial metabolism (Drews et al., 2010). Superoxide dismutase (*Sod*) converts reactive O_2_^−^ into less reactive hydrogen peroxide (H_2_O_2_). Catalase (*Cat*) and glutathione peroxide (Gpx) convert H_2_O_2_ into oxygen and water. Compared to the liver, expression levels of antioxidant molecules is much lower in pancreatic β-cells, with expression at 50% for Sod and only 5% for *Cat* and *Gpx* compared to liver, thus making them very susceptible to oxidative stress-induced damage (Tiedge et al., 1997). It is known that hyperglycemia and hyperlipidemia are linked to the elevation of ROS levels in β-cells. Although GSIS produces ROS via mitochondrial metabolism, chronic hyperglycemia in diabetes is linked to ROS accumulation, loss of mitochondrial membrane potential (Δψ_m_), and eventually β-cell apoptosis (Sivitz et al., 2010; J. Wang et al., 2017). In our studies, GYP treatment in BRIN-BD11 cells protected against H_2_O_2_ induced oxidative stress compared to untreated cells. Similar GYP protective effects against H_2_O_2_ have also been observed previously in retinal pigmental epithelial cells, vascular endothelial cells, and liver microsomes (Alhasani et al., 2018; Li et al., 1993). Thus, in β-cells, GYP may reduce the harmful effects of chronic ROS overproduction under metabolic stress and enhance β-cell function.

*Nrf2* is a master regulator of antioxidant gene transcription, which interacts with Kelch-like ECH- associated protein 1 (*Keap1*). Previous studies in db/db mice showed that *Nrf2* activation prevented β-cell oxidative damage and diabetes onset (Yagishita et al., 2014). In female Zucker diabetic rats, the *Nrf2-Keap1* pathway mediated β-cell self-repair after high fat diet-induced oxidative damage (Abebe et al., 2017). In isolated human islets, the *Nrf2* activator, dh404, increased antioxidant enzymes’ expression and decreased inflammatory mediators (Masuda et al., 2015). Previous studies in diabetic animal models and cell lines suggest that GYP elicits its cytoprotective effects by enhancing the *Nrf2* signaling pathway (Alhasani et al., 2018; Gao et al., 2016; Meng et al., 2014). In line with these reports, the current study in BRIN-BD11 cells produced a similar upregulation of *Nrf2* expression by GYP and its associated antioxidant genes *Sod1, Cat, Gpx1,* and *Ho1*. We also noticed a downregulation of *Nfkb1* and *Nfkb2* expression, which is associated with proinflammatory responses. *Erk1* (*Mapk3*) and *Erk2* (*Mapk1*) are dominantly expressed MAPKs in the pancreatic β-cells and are associated with cellular proliferation (Jiang et al., 2018). In rat INS-1 insulinoma cell lines, glucose and GLP-1 induced cell proliferation is linked to Ca^2+^ mediated *ERK1/2* activation (Arnette et al., 2003). In the current study, GYP enhanced *Erk1* mRNA expression at 24 and 72h while protein levels of total *Erk1/2* were unchanged (data not shown). It has previously been reported that in INS-1 cells, *Erk1/2* activation occurs within 30mins of exposure to high glucose or forskolin, indicating changes in *Erk1/2* occurs at a much earlier time scale than 24h or 72h, which we have used in the current study (Arnette et al., 2003). This could be the possible reason for unaltered protein levels of total *Erk1/2* by GYP at 24h / 72h.

Pancreatic/duodenal homeobox1 (*Pdx1*) and MAF BZIP Transcription Factor A (*MafA*) are the key transcription factors that bind to the insulin gene and promote its transcription. *Pdx1* plays a significant role in the development and function of β-cell, whereas *MafA* is necessary for GSIS (Melloul et al., 2002; Zhang et al., 2005). Post-translational modification of proinsulin by *Pcsk1* and *Pcsk2* produces bioactive insulin and C-peptide. Our findings have shown an increase in *Ins1* expression following GYP treatment, whereas *Pdx1* gene expression and protein level were reduced by 72 h GYP treatment. In contrast, *MafA* expression was unchanged along with *Pcsk1* and *Pcsk2* protein levels. Interestingly, *cMyc* levels are higher in juvenile islets but reduced in adult islets, and high levels of *cMyc* while enhancing β-cell proliferation, cause a decrease in *Pdx1* expression by binding to canonical binding sites upstream of *Pdx1*, but not *MafA* (Puri et al., 2018). This supports the idea that in BRIN-BD11 cells in the current study, GYP, while increasing *cMyc* and increasing cell proliferation, most probably does this at the expense of *Pdx1* expression and β-cell maturity.

Calmodulin is a ubiquitous protein associated with calcium-dependent activation of membrane-bound adenylyl cyclases (ACs) (Sharp et al., 1980). These ACs generate cAMP from ATP, and intracellular cAMP levels are further regulated by cyclic nucleotide phosphodiesterase (PDE) mediated degradation into 5’-AMP. cAMP, as a secondary messenger, exerts its functions predominantly through activating PKA, which has multiple cellular functions, including regulation of proliferation through phosphorylation of transcription factors such as *cMyc* (Padmanabhan et al., 2013) and *cJun* (Heinrich et al., 1997), regulation of mitogen-activated protein kinases like *Erk1/2* (Briaud et al., 2003) and *Jnk* (Hur, 2005), regulation of antioxidant response through *Nrf2* (Kulkarni et al., 2014) and anti-inflammatory response by inhibiting *Nfkb* activity (Takahashi et al., 2002). The current study shows that GYP enhances calmodulin (*Calm*) expression and downregulates *Pde3b* and *Pde4b* expression in BRIN-BD11 cells, indicating possible enhanced cAMP levels and perhaps downstream responses. However, cAMP and PKA pathways were not investigated in the current study as previous studies in isolated rat islets reported that GYP effect was mediated through PKA (Lokman et al., 2015). However, our results showed modulation of expression of genes both upstream and downstream of PKA suggesting an involvement of PKA activation in the beneficial effects of GYP in BRIN-BD11 cells.

## Conclusion

GYP enhanced Ca^2+^ uptake in BRIN-BD11 cells, which may be mediated by NCX reverse mode activation. GYP also enhanced insulin release irrespective of extracellular glucose concentration and showed cytoprotective effects against saturated free fatty acid palmitate, H_2_O_2_, cytokine cocktail, and enhanced antioxidant and pro-proliferative gene expression while downregulating proinflammatory response mediating genes. Further studies are required to confirm these findings in primary β-cells and human islets and to further establish and confirm the potential interaction with NCX channels involved in GYP action.

## Abbreviations

[Ca^2+^]_i_: Intracellular calcium
GYP: Gypenosides
GSIS: Glucose-stimulated insulin secretion

## Acknowledgments

We would like to thank Prof. Peter Flatt, Ulster University, for his kind gift of BRIN-BD11 cells.

